# High-Resolution Maps of Mouse Reference Populations

**DOI:** 10.1101/155507

**Authors:** Petr Simecek, Jiri Forejt, Robert W. Williams, Toshihiko Shiroishi, Toyoyuki Takada, Lu Lu, Thomas E. Johnson, Beth Bennett, Christian F. Deschepper, Marie-Pier Scott-Boyer, Gary Churchill, Fernando Pardo-Manuel de Villena

## Abstract

Genetic reference panels are widely used to map complex, quantitative traits in model organisms. We have generated new high-resolution genetic maps of 259 mouse inbred strains from recombinant inbred strain panels (C57BL/6J x DBA/2J, ILS/IbgTejJ x ISS/IbgTejJ, C57BL/6J x A/J) and chromosome substitution strain panels (C57BL/6J-Chr#<A/J>, C57BL/6J-Chr#<PWD/Ph>, C57BL/6JChr#<MSM/Ms>). We genotyped all samples using the Affymetrix Mouse Diversity Array with an average inter-marker spacing of 4.3kb. The new genetic maps provide increased precision in the localization of recombination breakpoints compared to the previous maps. Although the strains were presumed to be fully inbred, we found residual heterozygosity in 40% of individual mice from five of the six panels. We also identified *de novo* deletions and duplications, in homozygous or heterozygous state, ranging in size from 21kb to 8.4Mb. Almost two–thirds (46 out of 76) of these deletions overlap exons of protein coding genes and may have phenotypic consequences. Twenty-nine putative gene conversions were identified in the chromosome substitution strains. We find that gene conversions are more likely to occur in regions where the homologous chromosomes are more similar. The raw genotyping data and genetic maps of these strain panels are available at http://churchill-lab.jax.org/website/MDA.

## Introduction

The laboratory mouse is the most widely used mammalian model organism for biomedical research. Among the key advantages of mice are a well-annotated reference genome (Chinwalla *et al.* 2002), over one hundred strain-specific genome sequences (Keane *et al.* 2011), (Morgan *et al.* 2016), (CC Genomes, Genetics 2017), and many genetic reference populations, including multi-parent strain panels (Consortium 2012) and outbred stocks (Churchill *et al.* 2012), and strains carrying null alleles at most protein coding genes. There are dozens of readily available inbred strains that capture a wealth of genetic variants and display unique phenotypic characters (Beck *et al.* 2000), (Yang *et al.* 2011).

Genetic reference populations of mice include collections of strains that reassort a fixed set of genetic variants such as *chromosome substitution strain* (CSS) and *recombinant inbred strain* (RIS) panels. Chromosome substitution strains, also known as consomic strains, combine genomes of two founder inbred strains by substituting one chromosome pair from the *donor strain* into the genetic background of the *host strain* (Nadeau *et al.* 2012). The mouse genome is composed of 19 pairs of autosomal chromosomes, X and Y sex chromosomes, and a mitochondrial genome, thus a minimum of 22 strains could constitute a complete CSS panel. In some cases it has proven difficult to introgress a specific entire donor strain chromosome into the host background and the complete CSS panel may include partial chromosome substitutions and consists of more than 22 strains. RIS also combine genomes of two founder strains; they are derived from one or more generations of outcrossing followed by sibling mating to produce new inbred strains whose genomes are mosaics of the founder genomes (Williams *et al.* 2001). Both RIS and CSS panels have been successfully applied to the mapping of complex traits (Buchner and Nadeau 2015).

We have carried out high-density genotyping of three RIS panels C57BL/6J x DBA/2J (BXD), ILS/IbgTejJ x ISS/IbgTejJ (LXS), C57BL/6J x A/J (AXB/BXA) and three CSS panels C57BL/6J-Chr#<A/J> (B6.A), C57BL/6J-Chr#<PWD/Ph> (B6.PWD), C57BL/6J-Chr#<MSM/Ms> (B6.MSM) using the Affymetrix Mouse Diversity Array (MDA). The MDA includes approximately 623,000 probe sets that assay single nucleotide polymorphisms (SNPs) plus an additional 916,000 invariant genomic probes targeted to genetic deletions or duplications (Yang *et al.* 2009). These data add value to the strain panels by more precisely localizing the recombination breakpoints between founder strains. In addition they reveal some unexpected features in the genomes of individual strains.

## Materials and Methods

### Animals

We generated high-density genotype data for six mouse strain panels (Table 1): three panels of RIS and three panels of CSS. Mice for genotyping from five panels were available at the Jackson Laboratory (Bar Harbor, ME, USA) or from BXD colony at University of Tennessee Health Science Center (UTHSC); DNA samples from the sixth panel, B6.MSM CSS, were provided by T. Shiroishi (National Institute of Genetics, Japan). Unless stated otherwise, we genotyped one mouse per strain. Most strains are represented by a single male animal (255 males) but for four strains we genotyped an individual female (BXD14, BXD54, BXD59, BXD76). Samples were mainly from cases bred in 2008.

**Table 1.** : Overview of the six panels: a type, founder strains and a number of strains.

| Panel | Type | Founder strains |  | # strains |
| --- | --- | --- | --- | --- |
| AXB/BXA | RIS | C57BL/6J | A/J | 25 |
| LXS | RIS | ILS | ISS | 64 |
| BXD | RIS | C57BL/6J | DBA/2J | 91 |
| B6.A | CSS | C57BL/6J | A/J | 22 |
| B6.PWD | CSS | C57BL/6J | PWD/Ph | 28 |
| B6.MSM | CSS | C57BL/6J | MSM/Ms | 29 |

The AXB/BXA RIS panel (Nesbitt and Skamene 1984) was derived from intercrosses of the C57BL/6J (B or B6) and A/J (A) strains. Note that hereafter the dam is denoted first and the sire last. Thus the difference between AXB and BXA strains is the direction of the intercross mating that generated (AxB)F1s or (BxA)F1s, respectively. We genotyped 25 strains: AXB strains 1, 2, 4-6, 8, 10, 12, 13, 15, 18, 23, 24; and BXA strains 1, 2, 4, 11-14, 16, 17, 24-26.

The LXS RIS panel (Williams *et al.* 2004) was generated at the Institute for Behavioral Genetics, Bolder, CO from founder strains, Inbred Long-Sleep (L or ILS) and Inbred Short-Sleep (S or ISS). These founder strains were in turn derived as selection lines from a cross population with eight founder strains (A, AKR, BALB/c, C3H/Crgl/2, C57BL/Crgl, DBA/2, IS/Bi and RIII). We genotyped 64 strains: LXS 3, 5, 7-9, 13, 14, 16, 19, 22-26, 28, 32, 34-36, 39, 41-43, 46, 48-52, 56, 60, 62, 64, 66, 70, 72, 73, 75, 76, 78, 80, 84, 86, 87, 89, 90, 92-94, 96-103, 107, 110, 112, 114, 115, 122, 123.

The BXD RIS panel was derived from founder strains C57BL/6J (B or B6) and DBA/2J (D or D2) inbred mice in three epochs: epoch I, strains 1-32 (Taylor *et al.* 1975); epoch II, 33-42 (Taylor *et al.* 1999), and the epoch III advanced RIS 43-102 (Peirce *et al.* 2004b). The latter were outcrossed for multiple generations before inbreeding. We genotyped 91 strains: BXD 1, 2, 5, 6, 8, 9, 11-16, 18-25, 27-36, 38- 40, 42-45, 47-56, 59-71, 73-102 (note that the designation of several BXD strains have been modified as a result of the genotyping results described in the present study, and BXD103 is now known as BXD73b).

The B6.A CSS panel (Nadeau *et al.* 2000) consists of 22 strains derived from C57BL/6J (host) and A/J (donor) by J. Nadeau at Case Western Reserve University. The panel includes 19 autosomes, X and Y chromosomes, and the mitochondrial genome.

The B6.PWD CSS panel (Gregorova *et al.* 2008) consists of 28 strains derived from C57BL/6J (host) and PWD/Ph (donor) by J. Forejt at the Institute of Molecular Genetics AS CR in Prague, Czech Republic, covering all chromosomes and the mitochondrial genome. To improve reproductive fitness, chromosomes 10, 11 and X were split between three strains each carrying either the proximal (p), middle (m), or distal (d) portion of the respective chromosome.

The B6.MSM CSS panel (Takada *et al.* 2008) consists of 29 strains derived from C57BL/6J (host) and MSM/Ms (donor) by T. Shiroishi at National Institute of Genetics in Mishima, Japan covering all chromosomes. Chromosomes 2, 6, 7, 12, 13, and X were split between two strains each carrying either the centromeric (C) or telomeric (T) portion of the respective chromosome.

### Genotyping

Dna samples were prepared at the University of North Carolina according to the standard Affymetrix protocol and were hybridized on the Affymetrix Mouse Diversity Array (MDA) at the Jackson Laboratory as described previously in (Yang *et al.* 2009), (Didion *et al.* 2012). The MDA probes (NCBI37/mm9) were mapped to genomic positions in GRCM38/mm10 assembly. CEL files and updated mapping information are available at ftp://ftp.jax.org/petrs/MDA/raw_data/. We used the R software package MouseDivGeno (Didion *et al.* 2012) to extract intensities from CEL files, but for purposes of this study we developed a genotyping method that is based on the direct comparison of SNP probeset intensities between the sample and the founder strains of the corresponding panel. We selected the informative SNPs with intensity differences between founder strains for each panel (101,397 SNPs for AXB/BXA, 79,808 for LXS, 103,340 for BXD). Both selection of informative SNPs and SNP calls were probeset intensity based. For each strain and each SNP, the call can be either A (if the signal is close to the first founder), B (if the signal is close to the second founder), or N to represent “notA/notB”. We note that the N category includes both no-call and heterozygous genotypes and simply indicates that the intensity signal of the sample is far from both founder strains.

### Founder Haplotype Blocks

In order to define the haplotype blocks of founder genotypes with allowance for errors in individual SNP level genotype calls, we applied the Viterbi algorithm to smooth the genotyping. We used software implemented in the Hidden Markov Model (HMM) R package (Himmelmann 2010). We call the Viterbi algorithm iteratively: at each iteration we re-estimated the HMM transition probabilities based on the Viterbi reconstruction of haplotype blocks. The iterations are repeated until we reach the convergence (Juang and Rabiner 1990).

Genetic maps computed from RIS panels consist of intervals assigned to one of the founders and gaps that delimit the interval within which the inferred recombination event(s) have occurred. We refer to the latter as “recombination intervals”.

For RIS panels we compared our maps to those available at http://www.genenetwork.org.GeneNetwork.org provides two genotype files for the BXDs—a “classic” set (pre-2017) of genotypes that have been used in most mapping studies since 2005 (Shifman *et al.* 2006), and new consensus genotypes (2017) that include updated data for BXD43 through BXD220 that were collected November 2015 and processed using the GigaMUGA array (Morgan *et al.* 2016). In the current study we have compared MDA genotypes to the classic genotypes used through the end of 2016.

### Strain contamination

An RIS or CSS is considered to be contaminated if it carries a segment of genome that did not originate from one of the two founder strains. We developed an HMM to search for contamination. In contrast to our previous HMM analysis, here we select SNPs that were not informative (both founders have the same signal). In a contaminated region the signal of a given strain is expected to contain a higher proportion of SNPs that differ from both founder strains. To avoid only intervals covering three or more non-informative SNPs were reported.

### Copy number variants

To determine if any of the RIS or CSS strains carried copy number variations (CNVs) that differed from the copy number in the founder strains, we applied the *simpleCNV* function of the MouseDivGeno package (Didion *et al.* 2012). We accepted only those candidate CNV detections that had length >20kb and covered at least 10 IGP probes with *t*-statistic above 5 (p<1E-6).

### Gene conversions

Gene conversions are short tracts (<1kb) of nonreciprocal transfer of genetic information between two homologs that occurs during meiosis. In the case of RIS, it is difficult to distinguish gene conversion events from short haplotype blocks that are due to closely spaced recombination events that occurred in different meiosis. Therefore we restricted our attention to the CSS panels. We searched for single or small groups of adjacent SNPs that derive from the host genotype but occur on the donor chromosomes. We examined individual SNP intensities to identify those that are clearly derived from the host strain and are present in a region of donor strain haplotype.

### Sister strains

In a typical RIS panel the lineages that give rise to each RIS are independent and thus there should be no sharing of recombination events between strains. BXD strains from epoch III are an exception because they may share recombinations that arose in the outbreeding generations (Peirce *et al.* 2004a). Therefore, we excluded these strains from this analysis. We detected excess sharing of recombination junctions (Z-score>5.0) as an indicator that two strains are more similar than expected by chance.

## Results

**Global genotyping error** - defined as a percentage of informative SNPs discordant with the haplotype assigment - is typically below 1%, but it is higher for haplotype blocks of *M. m. musculus* (PWD) and *M. m. molossinus* (MSM) origin than for *M. m. domesticus* blocks (B6, A, D2) (Suppl. Figure 1). This is likely to be caused by polymorphisms in or near the oligonucleotide probe sequence or its flanking restriction sites (Didion *et al.* 2012). There are a few outlying strains with a higher error rate than other strains from the same panel (AXB1, BXD15, BXD25, BXD85, BXD65a (formerly known as BXD92), BXD93, B6.A#Chr7, B6.A#Chr10) likely due to low DNA quality or to processing of arrays.

**Residual heterozygosity** is present in some strains from each panel except for the AXB/BXA strains that appear to be fully inbred (Table 2). The detected heterozygous regions are an underestimate of percentage of segregating variation that is present in each strain because only a single animal per strain was genotyped. The presence of heterozygous strains in large RIS panels is not surprising. We estimated that in the absence of selection a RIS strain needs on average 24 generations of sib-mating to reach a heterozygosity rate below 1% and 36 generations to reach complete fixation. However, there is a significant variation in the number of generations required to achieve these landmarks (Broman 2005). For a panel of 22 strains (the size of a full CSS panel), 53 generations are required on average to achieve complete fixation for all its strains in the absence of selection.

**Table 2.** : Residual heterozygosity and CNV (deletion / extra copy) in the six panels.

| panel | number of strains | # strains with heterozygous segment |  | # strains with deletion |  | # strains with extra copy |  |
| --- | --- | --- | --- | --- | --- | --- | --- |
| **AXB** | 25 | 0 | 0% | 5 | 20% | 1 | 4% |
| **LXS** | 64 | 35 | 55% | 12 | 19% | 1 | 2% |
| **BXD, Epoch I** | 26 | 0 | 0% | 15 | 58% | 6 | 23% |
| **BXD, Epoch II** | 8 | 3 | 38% | 1 | 13% | 1 | 13% |
| **BXD, Epoch III** | 57 | 34 | 60% | 7 | 12% | 2 | 4% |
| **B6.A** | 22 | 3 | 14% | 2 | 9% | 2 | 9% |
| **B6.PWD** | 28 | 9 | 32% | 0 | 0% | 0 | 0% |
| **B6.MSM** | 29 | 2 | 7% | 12 | 41% | 1 | 3% |

### *De novo* deletions and duplications

We detected 64 *de novo* deletions and 14 *de novo* duplications, with lengths ranging from 21kb to 8.4Mb affecting 111 Ensembl genes (Suppl. Table 1). Table 2 summarizes frequency of strains with heterozygosity, deletions and duplications. We observe that longer time of inbreeding is associated with lower heterozygosity but more structural changes. This is seen most clearly by comparing different epochs of the BXD panel.

### High-density genotyping identifies unexpected haplotype blocks in CSS panels

We observe 27 haplotype blocks from the host strain in the proximal or distal regions of the donor chromosome across the three CSS panels. These events are undesirable but not unexpected due to the distribution of markers used for CSS development (Nadeau *et al.* 2000). We also observe strains in which a host haplotype block occurs in the middle of an introgressed donor chromosome or a donor haplotype block occurs in a host chromosome. We observed seven such events distributed across all three CSS panels. See Table 3 for details.

**Table 3.**
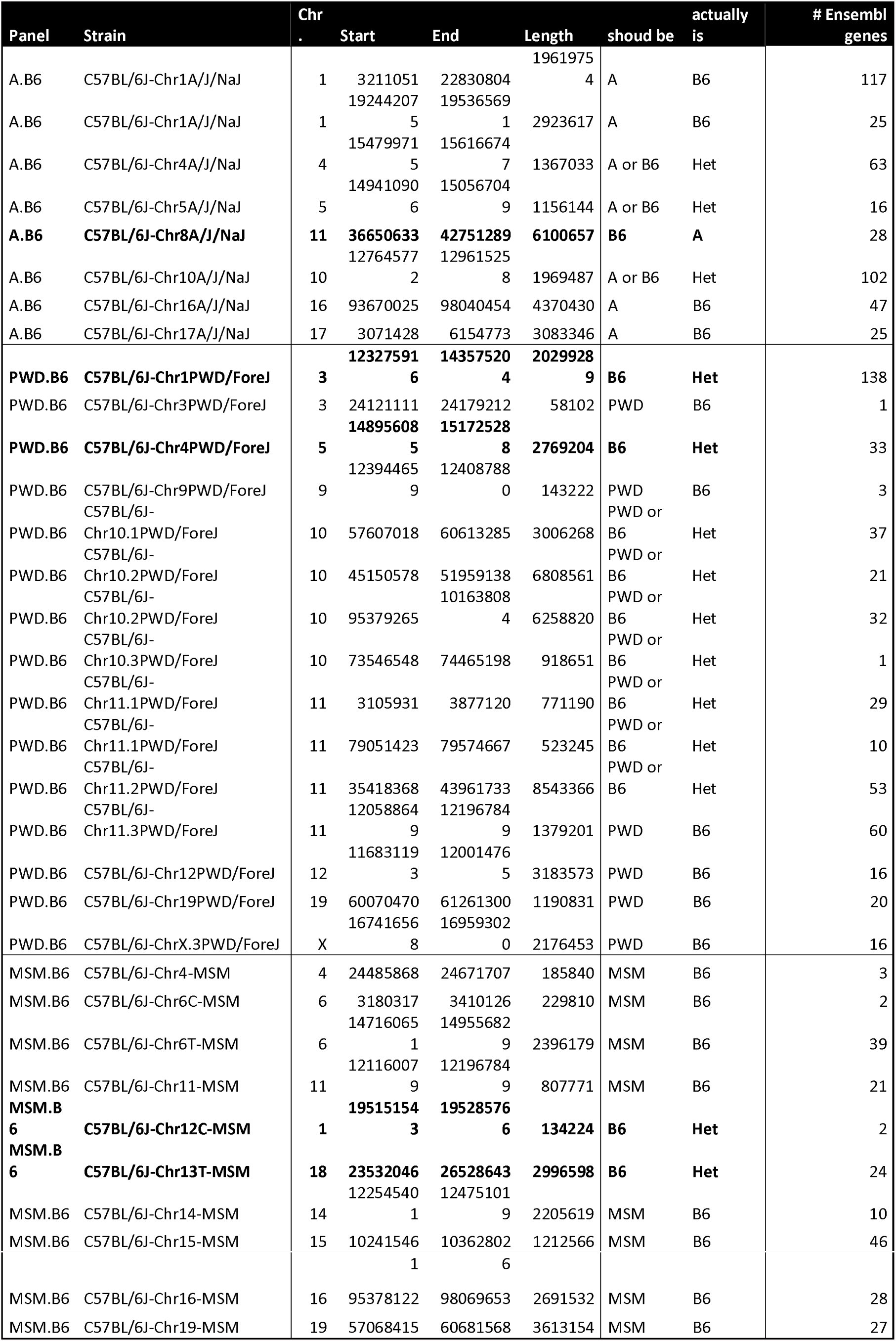
: Unexpected haplotype blocks in all CSS panels.

### High-density genotyping improves map accuracy in RIS panels

To validate our haplotype assignment and to estimate the level of improvement we compared our maps to the versions available at <u>www.genenetwork.org</u> (LXS, BXD) or provided by Institut de recherches cliniques de Montréal (AXB/BXA). There was a high concordance (99.8% LXS, 98.1% BXD, 99.5 ABX/BXA) between new and old maps for intervals that were in assigned to one of the founder in both maps. The new maps decreased the level of uncertainty measured as the sum of length of recombination intervals by 66% in the AXB/BXA panel, 41% in the BXD panel and 5% in the LXS panel. This improvement mirrors the increase in the number of informative markers: from 792 to 101,397 (AXB/BXA), from 3,796 to 103,341 (BXD), from 2,649 to 79,808 (LXS), respectively.

## Strain contamination in the AXB/BXA panel

An unexpected observation in AXB/BXA RIS panel, was the presence of six intervals that are not derived from either A or B6 inbred strains. Three chromosomes of AXB1 (x, y and z), two chromosomes of AXB2 and one chromosome of BXA1 (x and y) are affected. Based on comparison to genotypes from a large panel of inbred strains (Yang *et al.* 2011) we conclude that the contamination derived from a strain that is closely related to DBA/2J.

## Recombination rate

The distribution of the number of recombination events is similar across all panels (see Figure 1, Suppl. Table 2) with the exception of the advanced RIS BXD (epoch III) that has more recombination events per chromosome due to additional generations of outbreeding. The number of recombination events per strain ranges from 32 (BXD32) to 84 (BXA17) among the classical RIS and from 60 (BXD53) to 127 (BXD47) among the advanced BXD panel. These numbers of recombination events fall within the 95% prediction interval from simulations (using Python code from (Welsh and Mcmillan 2012)).

**Figure 1:**
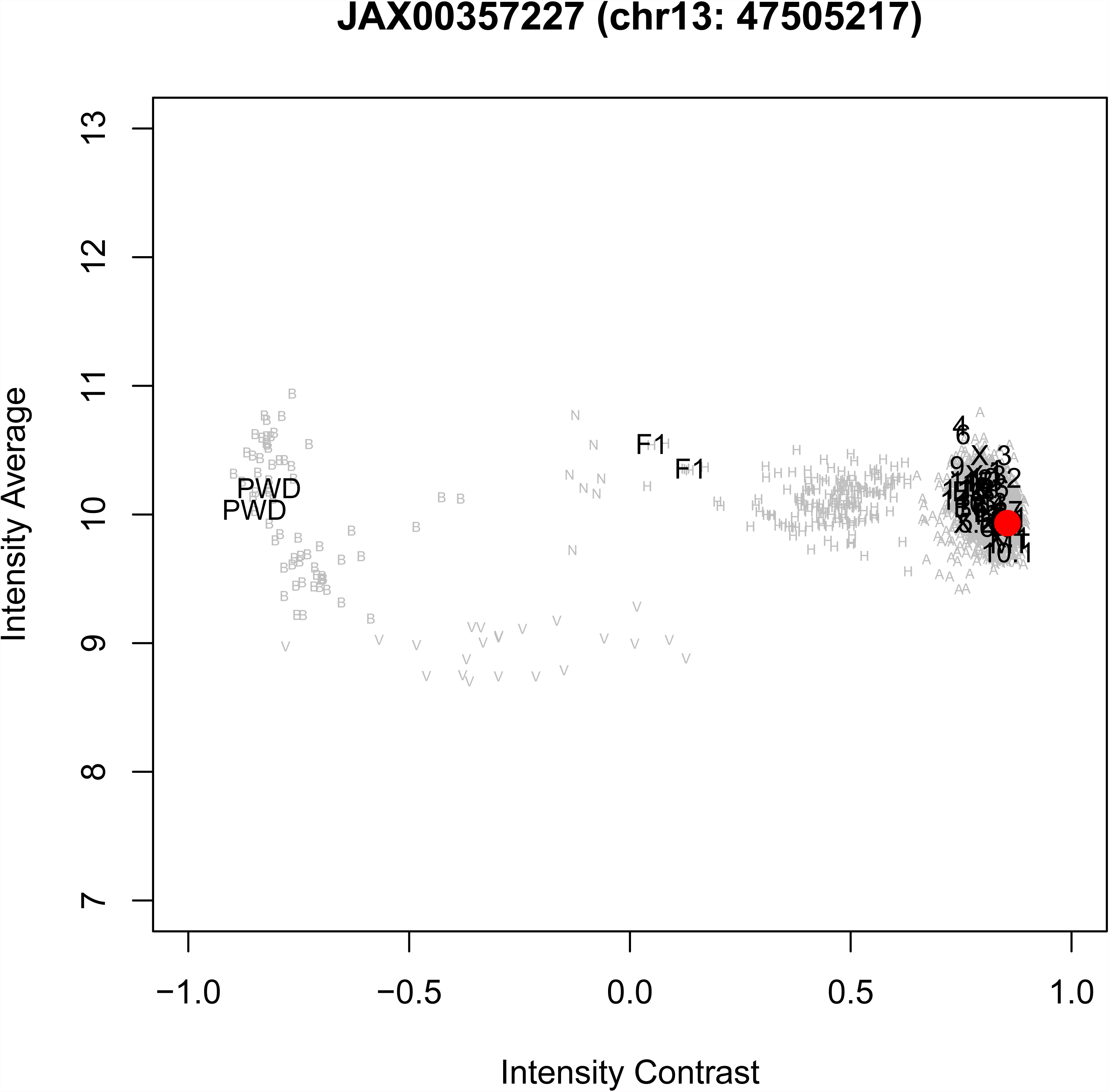
Number of founder haplotype blocks in RIS panels. The number of founder blocks for each strain is indicated as a point, with jitter for clarity. The boxplot indicates median and quartiles of each panel. Results for the BXD panel are broken down by three breeding epochs (I, II and III); the increased number of recombination event in epoch III refelects additional generation of outbreeding used in the derivation of these strains.

Most recombination events in the RIS panels are unique but some recombination intervals overlap and could result from independent recurrent events or from shared ancestry between RIS during the inbreeding process. The most frequently shared recombination event occurs in 8 out of 25 samples of the AXB RIS panel (Chr10: 66,730,215-67,348,211). Moreover, in 7 out of 8 cases (p=0.07) the polarity of the event is in the same direction: from B6 segment (proximal - 66730214 bp) to A/J segment (67348212 bp - distal). Additional shared recombination intervals are listed in Suppl. Table 3 and the recombination frequency is visualized in Suppl. Figure 3. Higher recombination rates observed in the distal region of chromosomes are expected (Liu *et al.* 2014).

## Sister strains

Sister strains are strains related by descent from incompletely inbred ancestors during the breeding process. They can be identified because they share a large number of recombination intervals with the same proximal to distal polarity of founder haplotypes. Not surprisingly, most of the sister strains are detected for the advanced BXD panel (6 pairs + 6 larger groups, totally comprising of 40 strains). However, two pairs of strains are present in the AXB and LXS panels, AXB6 - AXB12 and LXS94 - LXS107. These strains share more recombination intervals with the same founder strain polarity than expected by a chance (Figure 2).

**Figure 2:**
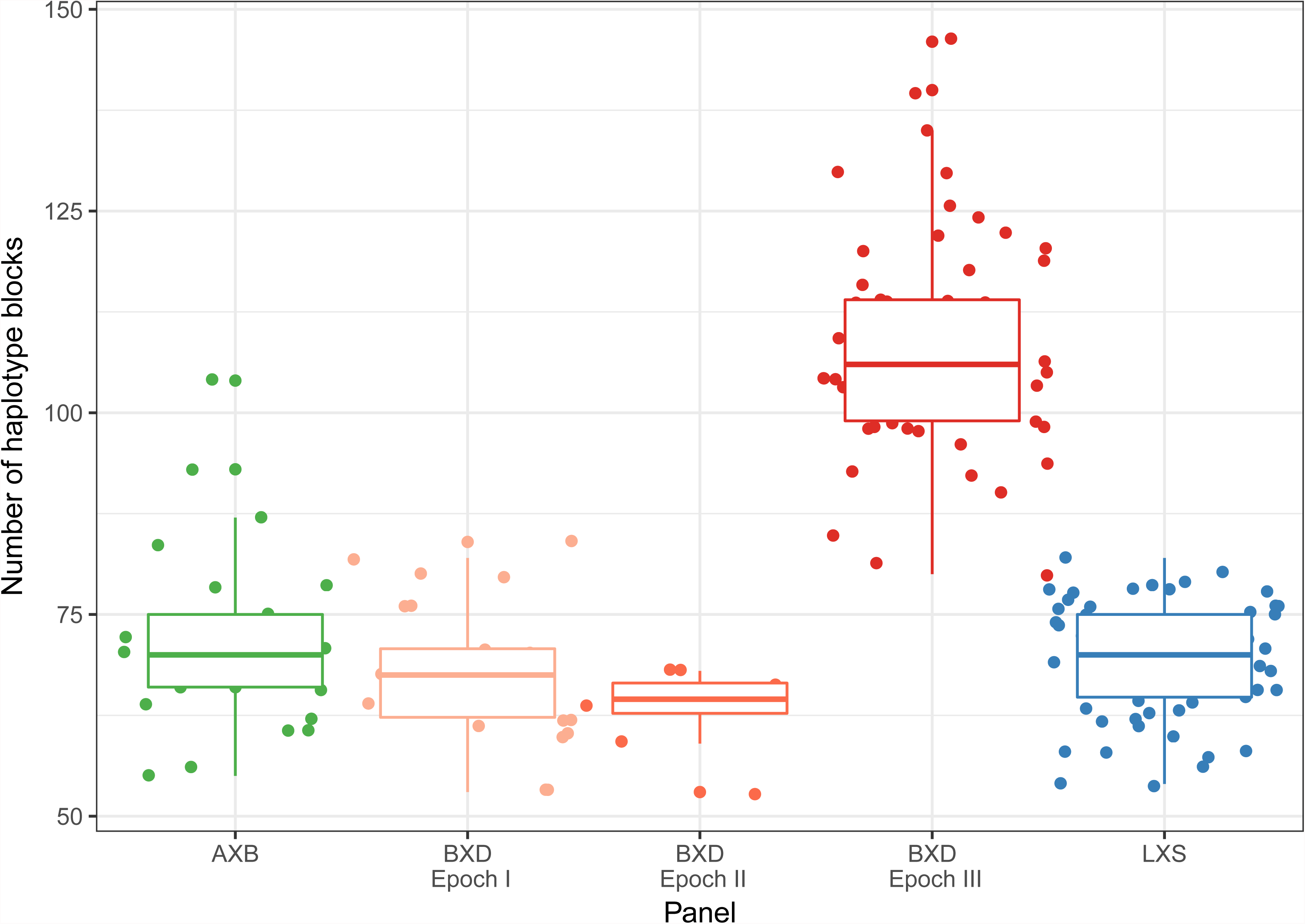
Sister strains in RIS panels. Side-by-side comparison of sister strains AXB6 vs AXB12 (red = B6, blue = A) (A) and LXS94 vs LXS107 (red = L, blue = S) (B) illustrates the extent of shared haplotype blocks.

## The MDA array detects short gene conversions in CSS panels

We searched for putative gene conversions in the introgressed donor chromosomes of CSS panels. We identified small regions typically spanning just one informative SNP, that have genotypes consistent with the host strain instead of the donor strain (Figure 3). In total, we identified 28 putative gene conversions: 17 in the B6.A CSS panel, 7 in the B6.PWD CSS panel and 4 in the B6.MSM CSS panel (Table 4).

**Figure 3:**
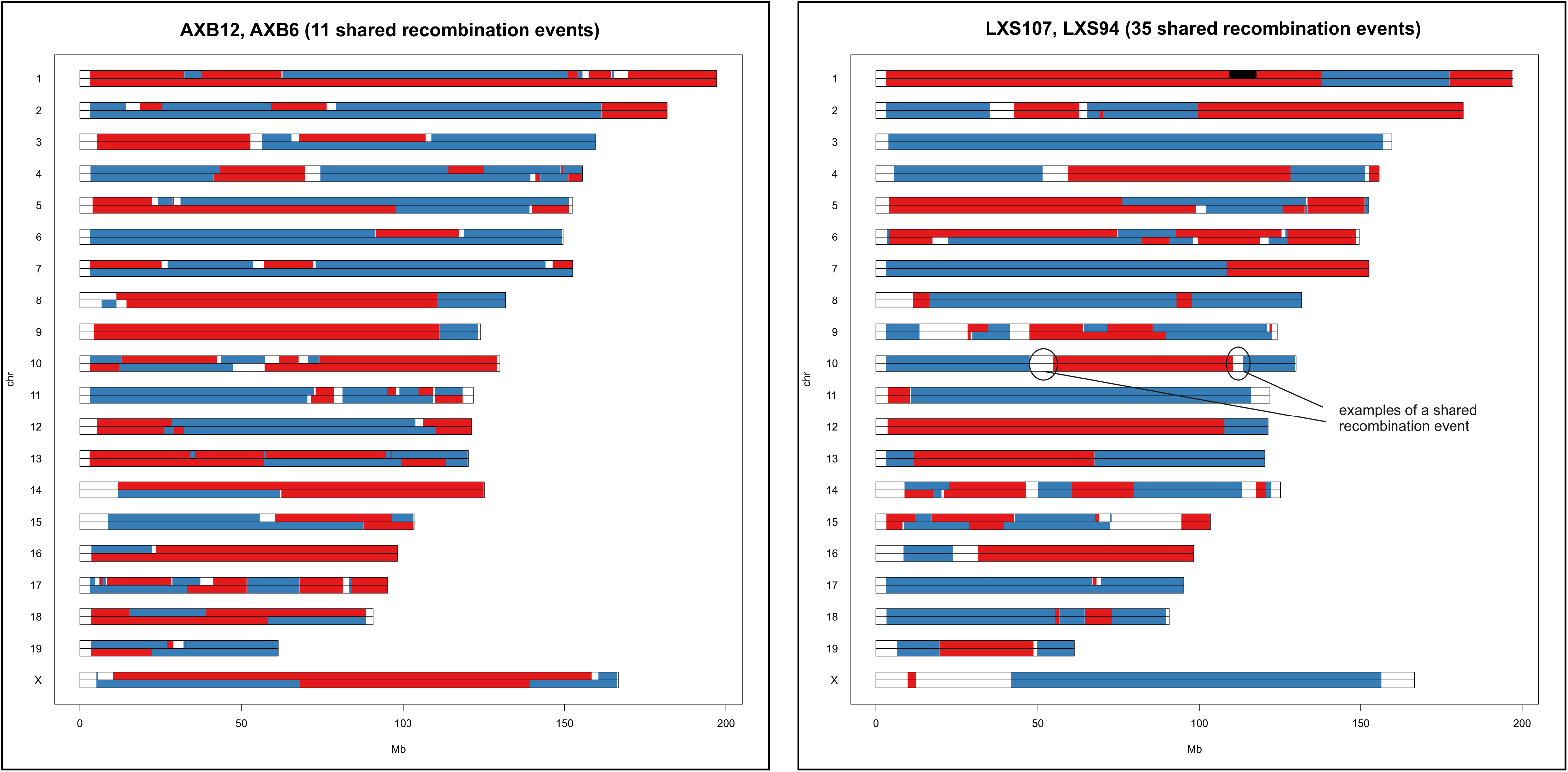
Gene conversion in a CSS strain. Strain B6.PWD13 has an unexpected founder genotype at marker JAX00357227 marker (Chr 13: 47,505,217 bp). Average and contrast signal intensities are plotted for all B6.PWD strains. Numbers indicate the CSS strains by substituted chromosome with B6.PWD13 is highlighted by the red circle. Also indicated on the plot are founder strains B6, and PWD and their F1 hybrids. The B.PWD13 data should be similar to PWD but it is actually close to B6 indicating a putative gene conversion. Grey letters indicate genotype calls for 1902 additional samples in the MDA database (A = first parent / B = second parent / H = heterozygous / V = vino / N = no call).

**Table 4:**
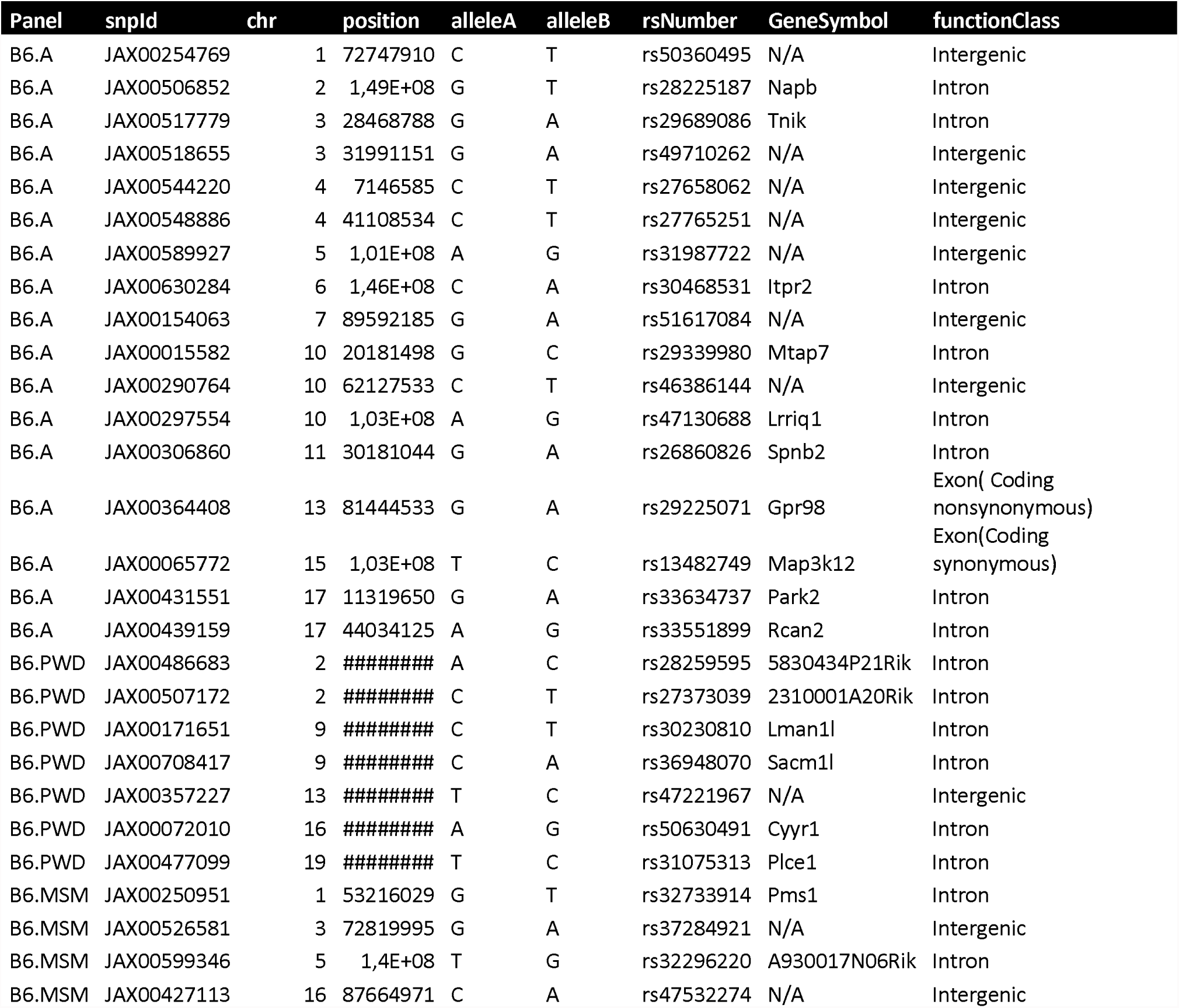
Short gene conversions in CSS panels.

## Online access to genetics maps and MDA genotypes

For easy access, we provide a compilation of Mouse Diversity Array data, annotation and supporting software at http://churchill-lab.jax.org/website/MDA. Resources to support our analysis of RIS and CSS strains include an online viewer where maps can be viewed and downloaded either as a list of intervals or as CSV files ready to be imported to the R/qtl package (Broman and Sen 2009). Source code for the viewer is also available on Github, https://github.com/simecek/RIS-map-viewer. Researchers interested in comparing those reference populations to genotypes of other mouse strains processed on MDA arrays can used the MDA viewer. The entire database consisting of 1,902 MDA arrays is available for download as SQLite database or as individual CEL files ftp://ftp.jax.org/petrs/MDA/.

## Discussion

We have characterized 180 RIS and 79 CSS strains from six popular and valuable resources and provided online access to these data. These panels were developed at different times and genotyped with lower density sets of markers. High-density genotyping with the number of informative SNPs, ranging between 79,000 and 257,000, provide maps with higher resolution. In this study we achieved a median spacing between informative markers 5.7 kb (AXB), 5.4 kb (BXD), 5.6 kb (LXS), 4.6 kb (B6.PWD) and 5.2 kb (B6.MSM), respectively. This enabled us to identify unusual features such as regions of residual heterozygosity, contamination by a non-founder strain and *de novo* structural variants. These genotyping arrays are part of 1902 samples processed on MDA platform that can be accessed from <u>http://churchill-lab.jax.org/website/MDA.</u>

Genetic reference panels are valuable, in part, because of the ability of generate animals with identical genomes in the number and timespan dictated by the researcher. Replication increases the accuracy of phenotype measurements (Belknap 1998) and allows for integration of data over space, time and environment. While it is convenient to think of all mice from an inbred strain as identical, we provide evidence that this view is not always warranted. Residual heterozygosity may be due to stochasticity in the inbreeding process or it may reflect biological constraints that prevent full inbreeding of a strain. Genetic drift operates in each of these populations and low-density genotyping in selected regions of the genome leaves room for undesired or unexpected surprises. In a typical CSS strain the average proportion of the donor genome present in other chromosomes is expected to be 0.2% (Nadeau 2000). Over our three CSS panels, the average length of unexpected genotype was 1.5 Mb. The length of intervals ranges (Table 3) from less than 1 Mb (1 gene) to 20 Mb (138 genes).

For gene conversions, whole genome sequencing of CSS panels (and RIS) will likely reveal more examples and provide better estimates of converted regions and their length. However, our results suggest that gene conversions are more probable in regions where founders’ genomes are very similar. We observe significantly more conversions on the B6.A panel that in the other two CSS panels (17 vs. 7 and 4, Fisher exact test, p=0.046) despite the fact that the number of informative markers is lower and therefore our ability to detect gene conversions reduced. Based on this result we hypothesize that gene conversions occur preferably in regions of low sequence diversity between homologous chromosomes. If that is true then they will have fewer genetic consequences due to lower chance to cause distinguishing polymorphism. Roughly, we estimate that 0.005% of the genome is affected by gene conversion (avg. # gene conversions / # informative SNPs = 28 / 3 / 200,000). The real number of gene conversions is likely to be higher because we were only able to identify gene conversions that overlap informative SNP probes in the array.

We found no evidence of allelic imbalance that has been observed in other species (Taudt *et al.* 2016). Nor did we detect any epistatic selection between founder strains or alleles with different subspecies origin. This is in sharp contrast with mouse multiparent populations such as the Collaborative Cross and Diversity Outbred (Chesler *et al.* 2016); (CC genomes 2017) and (Shorter 2017). Due to limited number of strains in mouse RI panels, we may have missed small distortions.

We observed an inverse relationship between residual heterozygosity and drift (Table 2). For a given panel, even 20 generations of inbreeding is not enough to fix all heterozygous regions. On the other hand, populations kept for many generations will accumulate SNPs, small indels, and structural variants in their genomes (Simecek *et al.* 2015) (CC genomes 2017). Strategies to reduce drift in breeding colonies have been developed, including the embryo cryopreservation program at The Jackson Laboratory (Taft *et al.* 2006). However, genetic drift can be also harnesed by geneticists to simplify and accelerate the identification of causal variants responsible for phenotypic differences between substrains (CC genomes 2017). These so call reduced complexity crosses are excellent examples of the potential benefits of genetic drift (Kumar *et al.* 2013).

## Acknowledgments

The work was supported by in part by NIH grants P50GM076468 (GAC), U19AI100625 (FPMV) and U01 AA016662 (RWW), a grant-in-aid for scientific research from Japan Society for the Promotion Science grant No. 17018033 (TS, TT), Canadian Institutes of Health Research MOP-93583 (CFD, MPSB) and by the Czech Science Foundation grant 16-01969S, LM2015040 and LQ1604 projects from the Ministry of Education, Youth and Sports of the Czech Republic (JF, PS). Genotyping of the BXD RIS was supported by the University of Tennessee Center for Integrative and Translational Science.

---

**Supplemental Figure 1**: Error rate. Each strain is plotted by two dots of different colors (one dot = one founder strain). A dot represents a percentage of markers contradicting the estimated founder strain.

**Supplemental Figure 2**: Percentage of genome attributed to the first or the second RIS founder strain (red = B6 or ILS, blue = A/J or D2 or ISS, green = heterozygous)

**Supplemental Figure 3**: Number of recombinations (smoothed by 10Mb window).

**Supplemental Table 1**: The list of RIS CNVs (deletion / extra copy).

**Supplemental Table 2**: Number of haplotype blocks (first founder / second founder / heterozygous) and the total number of recombinations.

**Supplemental Table 3**: The list of all recombination intervals (and the frequency of recombination).

